# Host-specific gene expression as a tool for introduction success in *Naupactus* parthenogenetic weevils

**DOI:** 10.1101/2021.02.23.432442

**Authors:** Ava Mackay-Smith, Mary Kate Dornon, Rosalind Lucier, Anna Okimoto, Flavia Mendonca de Sousa, Marcela Rodriguero, Viviana Confalonieri, Analia A. Lanteri, Andrea S. Sequeira

## Abstract

Food resource access can mediate establishment success in invasive species, and generalist herbivorous insects are thought to rely on mechanisms of transcriptional plasticity to respond to dietary variation. While asexually reproducing invasives typically have low genetic variation, the twofold reproductive capacity of asexual organisms is a marked advantage for colonization. We studied host-related transcriptional acclimation in parthenogenetic, invasive, and polyphagous weevils: *Naupactus cervinus* and *N. leucoloma*. We analyzed patterns of gene expression in three gene categories that can mediate weevil-host plant interactions through identification of suitable host plants, short-term acclimation to host plant defenses, and long-term adaptation to host plant defenses and their pathogens. This approach employed comparative transcriptomic methods to investigate differentially expressed host detection, detoxification, immune defense genes, and pathway-level gene set enrichment. Our results show that weevil gene expression responses can be host plant-specific, and that elements of that response can be transgenerational. Some host plant groups, such as legumes, appear to be more taxing as they elicit a complex gene expression response which is both strong in intensity and specific in identity. However, the weevil response to taxing host plants shares many differentially expressed genes with other stressful situations, such as host plant cultivation conditions and transition to novel host, suggesting that there is an evolutionarily favorable shared gene expression regime for responding to different types of stressful situations. Modulating gene expression in the absence of other avenues for phenotypic adaptation may be an important mechanism of successful colonization for these introduced insects.

## Introduction

### Invasiveness and diet breadth

Prevailing theories suggest that the majority of non-native species introduced to new habitats fail to establish due to travel stress, climate incompatibility, inadequate or inappropriate food resources, or small population size, among other factors (1). For species that do successfully establish, their potential to dynamically adapt phenotypic expression to environmental conditions (2) is thought to be correlated with establishment success (3). If the underlying biological causes behind invasion success were to be identified, global invasive control methods could be more targeted and efficient (4).

Food resource access can mediate invasive species establishment success (1). Herbivorous invasive insect species are on a continuum with regard to their dietary specialization (5). In general, species that feed on one or a few closely related plant species are considered to be monophagous “specialist” herbivores, whereas species that feed on more than one plant family are polyphagous “generalist” herbivores. Variation in species phenotype, including diet, can be caused by genetic and/or environmental factors (6). It has been proposed that a species that consumes varied host plants must account for differentiation between plants and develop an all-purpose phenotype (3), whereas a species that consumes one to a few plants can specifically optimize its usage: this set of evolutionary tradeoffs is often summarized as ‘jack of all trades, master of none’ (5). Other work has proposed that divergence in diet breadth is a byproduct of nonadaptive evolutionary forces such as drift (7). Data on gene expression in generalist herbivores supports the trade-off idea, finding that generalist herbivores have less fine-tuned gene regulation responding to different host plant diets, and broader patterns of gene regulation occur in generalists compared to specialists (8). In general, herbivorous insect generalists are thought to rely on transcriptional plasticity to respond to dietary variation (9, 10).

### Invasiveness and asexuality

The general-purpose genotype hypothesis (3) has been applied to invasive asexual organisms to explain their success. This hypothesis (as proposed by Lanteri and Normark, 1995) postulates that asexually reproducing species tend to have less strict habitat requirements, which allows wider spatial and environmental ranges compared to related sexual species. Asexually reproducing species require only one individual to start a population, which is advantageous for establishment and invasiveness (11). The twofold reproductive capacity of asexual organisms is a marked advantage for invasion (12, 13).

Within the weevil tribe Naupactini, several flightless species have been found to reproduce parthenogenetically (11). Reduced flight capacity has been hypothesized to be positively related to parthenogenetic species colonization in heterogeneous landscapes (13). Furthermore, flightlessness and obligate parthenogenesis have been linked to extreme polyphagy in successfully invasive insects (12).

*Naupactus* weevils are a taxonomic group of approximately 170 species of medium-sized weevils, covering a native geographic range between Mexico and Argentina (14–16). The asexual weevil species *Naupactus cervinus* and *N. leucoloma* reproduce via apomictic parthenogenesis, in which offspring are produced from unfertilized, diploid egg cells. Parthenogenetic species are thought to establish successfully due to their ability to preserve successful genotypes via clonal reproduction, as beneficial gene relationships are preserved under extreme linkage disequilibrium (17).

Fuller’s rose weevil, *Naupactus cervinus*, is a highly polyphagous species (12, 18). Native to South America, it has successfully established invasive populations in many countries via commercial trade, including the United States and Australia (19). The white-fringed weevil, *Naupactus leucoloma*, is also parthenogenetic, invasive, and highly polyphagous (20). Native to central and northern Argentina, southern Brazil and Uruguay, this species has successfully established populations in Chile, Peru, Australia, New Zealand, South Africa, and the United States (21). Most damage to crops and other host plants by both weevil species is caused by larvae feeding on roots, while the damage caused by the leaf feeding adults is usually less significant (22).

Even in the absence of genetic variation, parthenogenetic species can still become successfully established invasive species. In that case, what kinds of genetic and/or transcriptional adaptation and acclimation do these parthenogenetic species employ to acclimate to a new environment?

### Differential gene expression in targeted gene categories

A comparative transcriptomic approach was employed to measure phenotypic variation of *Naupactus* weevils in response to host plant type, specifically focusing on differential expression of genes that mediate functions that may impact invasion success. We also explored transgenerational changes in gene expression, given that transgenerational epigenetic modifications have been found to impact the expression of fundamental survival traits (i.e. lifespan and age at maturity) (23, 24).

One well-documented group of genes important for finding suitable host plant species in herbivorous insects are host detection genes associated with olfaction and taste, such as odorant-binding proteins (25–27). Another key functionality important for herbivore adaptation is that of detoxification and neutralization of plant secondary compounds; differential regulation of detoxification genes has been correlated with successfully feeding on new host plants (28). Detoxification genes may form the short-term first line of defense for herbivorous insects introduced to a new host (29). Moreover, detoxification of host plant defenses may continue to be a challenge given that a generalist’s longer-term response to a new host has been shown to include three times more differentially expressed genes related to detoxification (30). Gene pathways known to be involved in the detoxification response of herbivorous insects include *cytochrome p450*, *gluthathione-S-transferases*, *UDP-glycosyltransferases*, *carboxylesterases*, *ABC transporters*, and *glutathione peroxidases* (26,31,32).

Generalist herbivores feeding on a variety of plants are also often exposed to a wider range of pathogens and toxins, which drives a stronger selective pressure on generalists’ immune systems (33). Host plants of different nutritional qualities and defense capacities could alter herbivore immune defense response in a species-specific manner (33, 34), or generalists may have evolved general immune defense upregulation mechanisms that do not vary between hosts (35). Alternatively, resource investment in immune defense mechanisms could decrease as introduced invertebrates move away from their co-adapted pathogens (4). According to the enemy release hypothesis (36), individuals in a new environment will reallocate resources associated with immune defense towards growth and reproduction.

Our analysis therefore includes five groups of contrasts as detailed below: two between weevils feeding on hosts from different plant families (Legume vs. Other, Legume vs. Citrus); one between weevils feeding on hosts under different cultivation conditions (Conventional vs Organic); one between weevils feeding on host plants within the same host plant family (including members from Rutaceae, Fabaceae, and Asteraceae); and finally, one between weevils feeding continuously on one host plant versus weevils that have been transferred onto a previously un-encountered, but edible host. In those contrasts we will explore differential regulation of genes related to olfaction and chemosensory cues, those related to detoxification of host plant secondary compounds, and those related to immune system response genes. We will explore differences in the number of upregulated genes, the intensity of the increased expression (measured in fold change and other indexes of differential expression), and numbers of uniquely differentially expressed genes in all three gene categories in both immature and adult tissues.

### Predictions for expression patterns in weevils feeding on Legume hosts vs. other (non-legume) hosts

As potential host plants, legumes harbor a high diversity of defensive secondary metabolites, including alkaloids, amines, cyanogenic glucosides, and non-nitrogen-based compounds such as phenolics and terpenoids (37). Cyanogenic glucosides in particular are lethal to most herbivores, as they can disrupt cellular respiration and effectively shut down cellular functionality. Nitrogen-based defensive compounds are fairly unique to Fabaceae due to their association with nitrogen-fixing rhizobia. High levels of nitrogen in host plants are preferred by insect herbivores (38), because insects cannot produce their own nitrogen and must derive nitrogen nutritionally (9, 25). Previous studies indicate that herbivorous insects perform best on plants with high levels of rhizobial interactions (38). Because these legume-specific chemical defenses are damaging to herbivorous insects, there is a strong evolutionary pressure on legume-feeding species to develop adaptive mechanisms by which they can effectively break down these nitrogen-based defensive compounds (25). In *Naupactus* specifically, *N. cervinus* larvae performed better on a legume host (18), and *N. leucoloma* has been shown to prefer legume species (39). We predicted that when comparing differentially expressed genes between *N. cervinus* and *N. leucoloma* weevils feeding on legume host plants versus other (non-legume) host plants, there will be more differential regulation of genes in the three targeted categories in weevils feeding on legume host plants in both adult and immature tissues.

### Predictions for expression patterns in weevils feeding on Legume hosts vs. Citrus hosts

Citrus (family Rutaceae: subfamily Citrinae) also produce a variety of defensive secondary metabolite compounds, such as limonoids, flavonoids, alkaloids, carotenoids, and phenol acids (40). As some of these defensive compounds are unique to citrus, successful citrus herbivore species must have some counteracting or defensive mechanisms to allow them to survive. Despite the systemic nature of many citrus species’ defense responses, some of the strongest chemical defenses produced by citrus, such as limonene, occur in the fruit itself, which *Naupactus* does not consume (ex. (41)). Because *Naupactus* larvae feed on root tissue while adults feed on leaf tissue, it is likely that the secondary metabolites produced by legumes will be more deleterious to *Naupactus* weevils than those produced by citrus. We predicted that when comparing differentially expressed genes between *N. cervinus* weevils feeding on legume host plants versus citrus host plants, there will be more differential regulation of genes in the three targeted categories in weevils feeding on legume host plants in both adults and immature tissues.

### Predictions for expression patterns in weevils feeding on organically grown vs. conventionally grown oranges

There is inconsistent evidence regarding the effects of organic versus conventional farming techniques on agricultural pest burdens. Some research proposes that generalist diets predispose herbivorous insects towards evolving effective insecticide resistance, making feeding on conventional hosts less costly (7). Regardless of herbivore diet breadth, the assumption is that applying insecticides to host plants will make insect feeding more difficult, and conversely, reducing chemical insecticide usage on plants will increase the pest burden (42). However, no significant correlation was found between pest damage and farming management approaches for garden tomatoes (42); it is possible that organically grown plants *not* exposed to insecticides are capable of synthesizing their own chemical defenses. The addition of insecticides to a conventionally raised plant may interfere with the natural defense response of the plant, and an organically raised plant may be able to upregulate its defensive response in ways that conventionally raised plants cannot. We predicted that when comparing differentially expressed genes between weevils feeding on organically treated host plants versus conventionally treated host plants, there will be more differential regulation of genes in the three targeted categories in weevils feeding on organically cultivated host plants in both adults and immature tissues.

### Predictions for expression patterns in weevils feeding on different host plants within the same host plant family

If it is true that legume and citrus hosts are more resource-taxing to herbivores compared to other hosts, it could be expected that herbivores that feed on highly chemically defended species will have more species-specific transcriptional responses, and that the weevils consuming these host plants have acclimated to these defenses.

Because of this acclimation, we predicted larger numbers of unique expression patterns between weevils feeding on citrus members (Rutaceae), and between those feeding on legume members (Fabaceae), than between those feeding on members of a non-citrus, non-legume group (Asteraceae), even though the degrees of phylogenetic relatedness between host plants within each family are not equivalent. Furthermore, there will be more differential regulation of genes in the three targeted categories in weevils feeding on legume and citrus host plant family members relative to those from the non-citrus, non-legume host plant family comparisons.

### Predictions for expression patterns in weevils feeding on their natal host plant vs. a novel host plant

In polyphagous herbivores that can consume several host plants, a shift from consuming one host plant to a different host plant has been previously associated with high transcriptional responses (8,9,29). Although patterns of transcriptional response to short-term host plant switching are characterized by highly specific gene responses, these responses occur within a small number of gene families, indicating the potential for common pathways of host plant acclimation and adaptation in generalist arthropods (29).

When comparing differentially expressed genes between *N. cervinus* weevils feeding on their natal host plant versus those feeding on a novel host plant, we predicted that that there will be more differential regulation of genes in the three targeted categories in weevils feeding on the novel host in both adults and immature tissues.

### Exploration of global expression patterns in all host plants and experimental contrasts

It is entirely possible that important aspects of weevil acclimation and/or adaptation to feeding on resource-taxing host plants, or on novel hosts, may involve differential regulation of genes beyond the three targeted gene categories of detection, detoxification and immune response. For example, a plastic response, as measured by a wider array of upregulated gene sets, was recorded in milkweed aphids feeding on novel host plants (43), and specific gene expression response trajectories were elicited in response to different sugar-mimic alkaloids in silk moths (44).

In insects, developmental gene networks are well-known and have been profiled in several species (45); thus, it is not implausible to hypothesize that other tightly synchronized gene networks might exist. Metabolic pathways have also been found to be key in herbivore response to host plant defenses (46). We use a global gene set enrichment approach to elucidate overall patterns of expression by gene family. Together with the observed patterns in the three targeted gene categories, the goal of these analyses is to understand the role of host plant acclimation and adaptation in introduced species.

We sought to profile the transcriptome of successfully invasive, but paradoxically asexual, insects, and determine how life stage, host plant, and environmental conditions affect gene regulation in these species. We have successfully established that gene expression response of weevils can be specific to particular host plants, and that elements of that response can be transgenerational. We have gained understanding of how some host plants are more taxing to weevils eliciting strong and specific gene expression response. However, we also found commonalities to the response of taxing host plants and other stressful situations such as host plant cultivation conditions and/or a transition to a novel host.

## RESULTS

### Weevils display host-specific gene expression responses in the three targeted gene categories

#### Legume host plants generate large transcriptional responses

For both *N. cervinus* and *N. leucoloma*, there were significantly more upregulated host detection (HD), detoxification (DTX), and immune defense (IM) genes in weevils feeding on legume host plants (Fig 1i). In legume-feeding weevils, odorant-binding proteins were the most numerous HD genes overexpressed; *cytochrome P450* type genes were the most numerous DTX genes overexpressed; and serine proteases and proteinases were the most numerous IM genes overexpressed (Fig 1i). Furthermore, the differences in the numbers of upregulated genes between host plants in all three gene categories were significantly impacted by the different functional groups within each category, displaying significant interactions between Host:Functional Gene Group (Table 1). However, there was no discernible effect of tissue type for any of the three target gene categories, as evidenced by non-significant Host:Tissue interactions (Table 1). Finally, when exploring the interaction of tissue effects and functional group effects on differentially expressed gene (DEG) differences between host plants, we found that both factors significantly impacted the expression of DTX genes only in *N. cervinus*.

**Fig 1.**
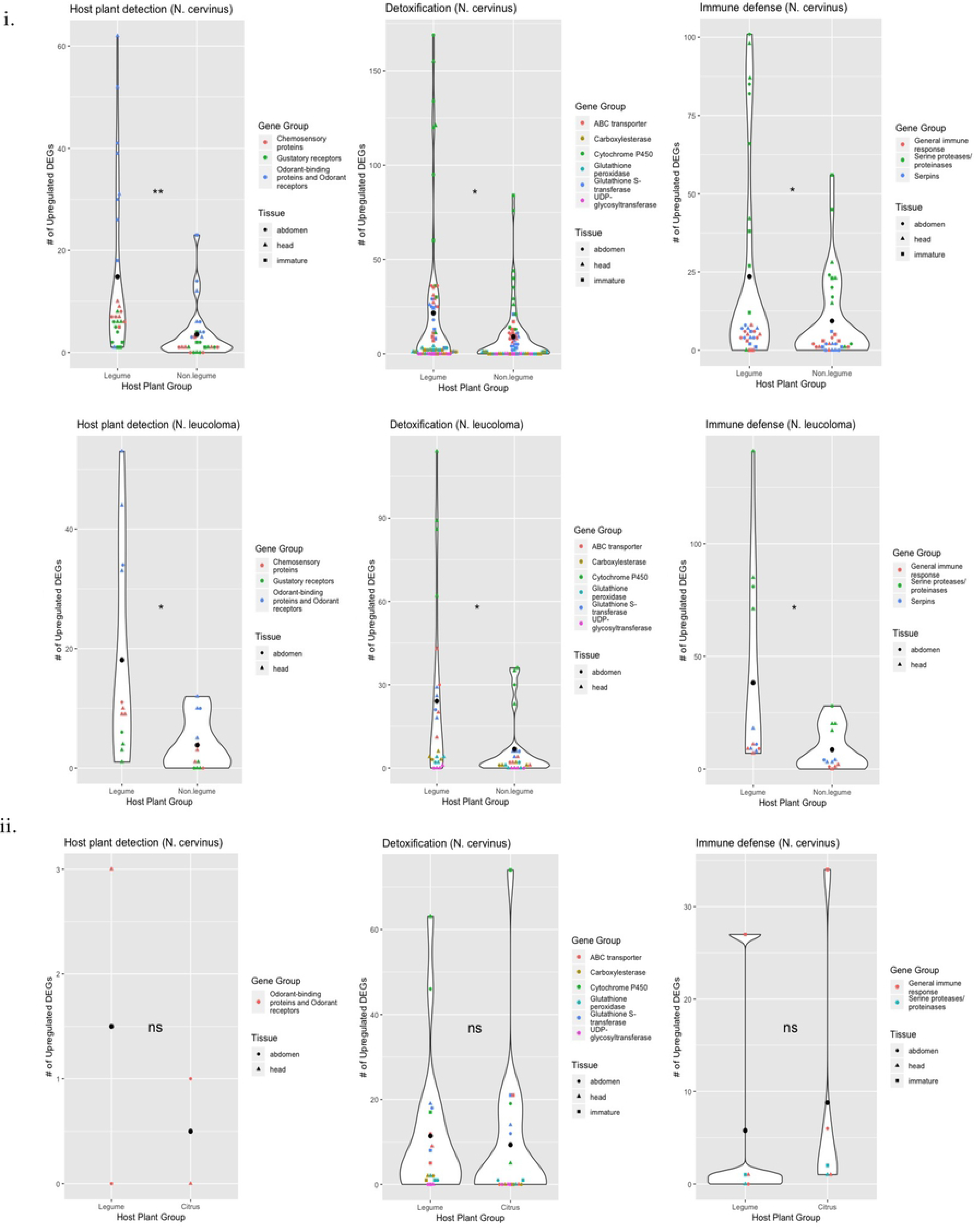
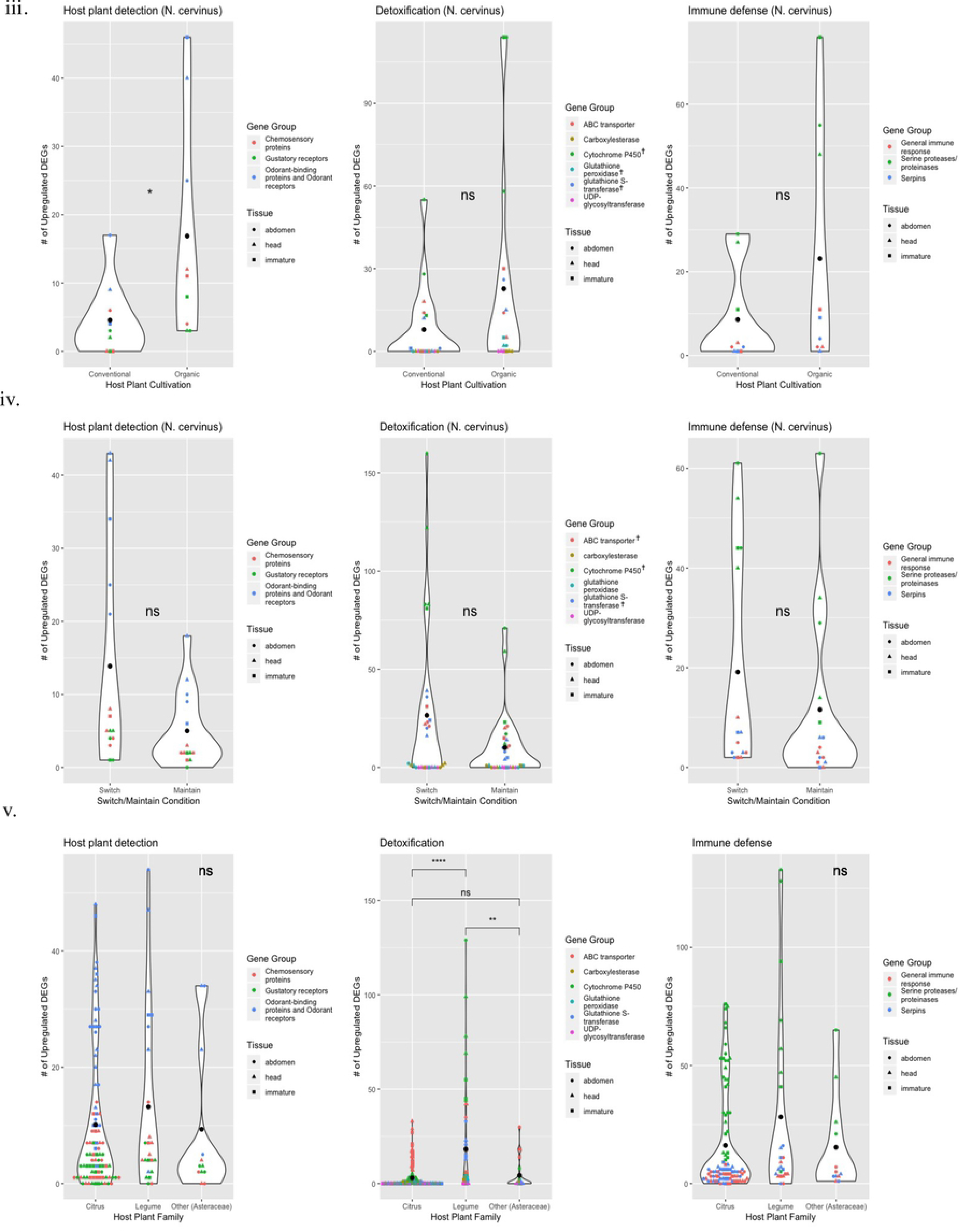
Number of differentially upregulated genes in three targeted gene categories from weevil species feeding on different host plants or in different experimental conditions. Categories analyzed include genes related to host plant detection (HD), host plant detoxification (DTX) and immune defense (IM) when comparing: (i) weevils feeding on Legume vs. Other in *N. cervinus* and *N. leucoloma*; (ii) *N. cervinus* weevils feeding on Legume vs. Citrus; (iii) *N. cervinus* weevils feeding on oranges grown under Conventional vs. Organic farming methods; (iv) *N. cervinus* weevils maintained on the natal host plant or switched from that host to a novel host - Switch vs. Maintain; (v) weevils feeding within the same host plant family: Citrus (Rutaceae: Citrinae), Legume (Fabaceae), or Other (Asteraceae) host plants. Each point represents a separate pairwise comparison in the set; i.e. one triangle represents the number of DEGs from a head tissue comparison that belongs in the group ‘Legume vs. Citrus’.

**Table 1.** Summary of comparisons of the numbers of differentially upregulated genes in three gene categories for weevil species feeding on different host plants or experimental conditions.

| Host plant contrasts | Species | Functional Gene Group | Differences between #s of upregulated genes when feeding on different host plants | Interaction |  |  |
| --- | --- | --- | --- | --- | --- | --- |
|  |  |  |  | Host: Tissue | Host: Functional Gene Group | Tissue: Functional Gene Group |
| Legume vs. Other | *N. cervinus* | HD | <0.05* | NS | <0.05* | NS |
|  |  | DTX | <0.05* | NS | <0.01** | <0.01** |
|  |  | IM | <0.05* | NS | <0.01** | NS |
|  | *N. leucoloma* | HD | <0.05* | NS | <0.01** | NS |
|  |  | DTX | <0.05* | NS | <0.01** | NS |
|  |  | IM | <0.05* | NS | <0.01** | NS |
| Legume vs. Citrus | *N. cervinus* | HD | NS | NS | NS | <0.05* |
|  |  | DTX | NS | NS | <0.01** | NS |
|  |  | IM | NS | NS | NS | <0.01** |
| Conventional vs. Organic | *N. cervinus* | HD | <0.05* | NS | <0.05* | NS |
|  |  | DTX | NS ‡ | NS | <0.01** | NS |
|  |  | IM | NS | NS | <0.01** | NS |
| Within-host Family (Citrus, Legume, Other) | *N. cervinus, N. leucoloma* | HD | NS | NS † | NS | NS |
|  |  | DTX | <0.05* (C-L; L-O);<br>NS (C-O) | <0.01**† | <0.01** | <0.05* |
|  |  | IM | NS | NS † | <0.01** | <0.01** |
| Switch vs. Maintain | *N. cervinus* | HD | NS | NS | <0.01** | NS |
|  |  | DTX | <0.05*‡ | NS | <0.01** | NS |
|  |  | IM | NS | NS | <0.01** | NS |

Results are displayed by prediction, species and gene category (HD: host detection; DTX: host detoxification and IM: immune defense). *P*-values displayed indicate significance for contrasts between host plants (Wilcoxon signed-rank test). ANOVA interaction terms were calculated between host effects and those of tissue (Host:Tissue) and functional gene groups within each gene category (Host:Functional
Gene Group) and between tissues and functional gene groups (Tissue:Functional Gene Group) (Robust rank-based ANOVA in package `Rfit`, derived from Hettmansperger and McKean (2010) and Hocking (1985). † indicates Kruskal-Wallis. ‡ indicates several individually significant functional gene groups. \*, \*\* indicate significance at $p=0.05$ and $p=0.01$ , respectively.

Heatmaps revealed not only the number of upregulated genes, but also the intensity of expression for the DEGs that were differentially regulated. For *N. cervinus*, there was strong expression intensity for both DTX and IM genes in legume-feeding weevil abdomen samples (Fig 2). Immature tissue also showed upregulation of DTX and IM genes, but expression intensity was high from both legume-feeding and other-feeding parents (Fig 2). For *N. leucoloma*, a clearer pattern was visualized wherein HD, DTX, and IM genes all had higher expression intensities in head tissue from legume-feeding weevils, but all three gene classes had higher expression intensities in abdominal tissue from other-feeding weevils.

**Fig 2.**
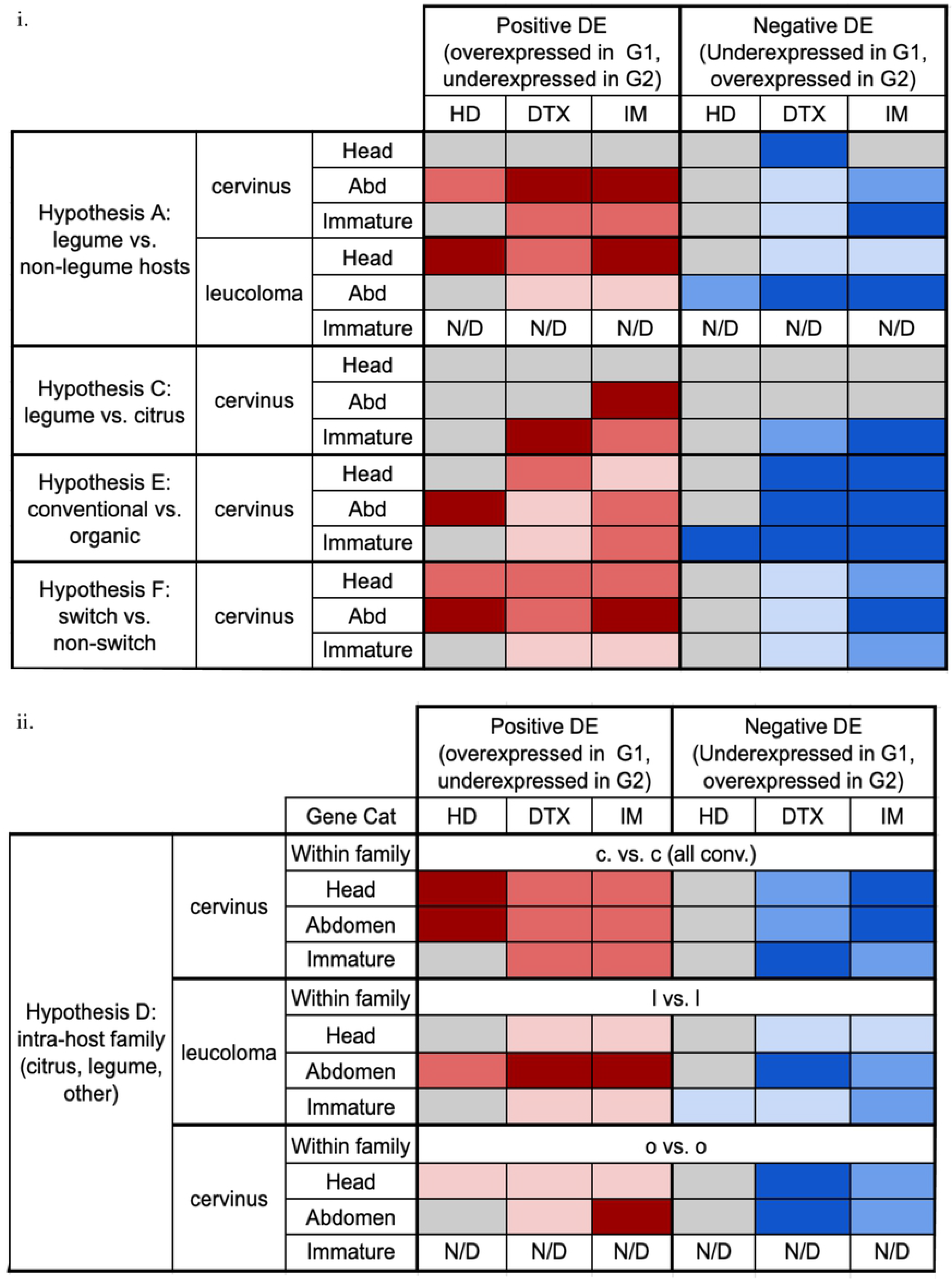
Composite heatmap showing expression intensity of significantly up- and downregulated genes in three gene categories including all available tissue types for weevils feeding on different host plants or in different experimental conditions. Results are displayed by prediction, species and tissue for each gene category (host detection (HD), host detoxification (DTX) and immune defense (IM)) and each direction of expression. (i) contrasts when feeding on different plant families, farming methods and experimental conditions, (ii) feeding within the same host plant family: Citrus (Rutaceae: Citrinae) (c vs. c); Legume (Fabaceae) (l vs. l); and Other (Asteraceae) (o vs. o). Shades of red indicate upregulation in Group 1 while shades of blue indicate upregulation in Group 2.

#### Citrus host plants elicit similar numbers of herbivore DEGs relative to legumes, with interesting patterns in pre-feeding immatures

Very few HD and IM genes made the cutoff criteria in these comparisons, and the number of upregulated genes are non-significantly different between host plants. A larger number of DTX genes made the cut-off criteria but neither the aggregate data, nor the data analyzed by gene, yielded significant differences between host plants (Fig 1ii). Despite the non-significant effects of host plant in all gene categories, we found that the differences in the numbers of upregulated DTX genes between host plants was significantly impacted by the different functional groups within that category (with a significant Host:Functional Gene Group interaction). Again, there was no discernible effect of sampled tissue type, as evidenced by a non-significant Host:Tissue interaction (Table 1). Interestingly, the interaction of tissue effects and functional gene group effects on the DEG differences between host plants was significant for the expression of HD genes, but not for DTX genes in *N. cervinus*.

Heatmaps revealed that the weighted median expression intensity of DTX and IM genes in immature tissue with legume-feeding as well as citrus-feeding parents was quite high, indicating that both host conditions elicited differential expression in a similar number of genes of different identities. There was also strong expression of IM genes in abdominal tissues of legume-feeding adults (Fig 2i).

#### Organically raised host plants elicit higher numbers of some herbivore host detection and detoxification genes relative to conventionally raised hosts

As expected, there was a significantly higher number of HD-related olfaction and chemosensory genes upregulated in weevils feeding on organically cultivated host plants (Table 1, Fig 1iii). There was no significant difference between number of DTX genes when analyzed as an aggregate, but weevils feeding on organically grown oranges showed significantly larger numbers of upregulated genes in three specific DTX gene functional groups (*cytochrome p450*, *glutathione peroxidase* and *glutathione S-transferase*). Despite non-significant host effects on the number of upregulated genes in the IM gene category, there was a significant interaction in the Host:Functional group for all three gene categories (Table 1).

Heatmaps revealed a high intensity of expression for HD genes in immature tissue from larvae produced by weevils feeding on organically raised oranges, as well as in abdominal tissue from weevils feeding on conventional oranges (Fig 2i). DTX genes had a higher median expression intensity in organically-feeding weevils compared to conventionally-feeding weevils in abdominal and immature tissues, although this did not hold true for all tissues; higher expression intensity was detected in head tissue of conventionally-feeding weevils.

#### Within host plant family contrasts revealed stronger detoxification response in weevils feeding on different legumes

Quantities of both HD and IM genes were not significantly different in contrasts within each host plant family (citrus, legumes, and asters), although there was a slightly higher number of both IM and HD genes upregulated in intra-legume comparisons (Fig 1v). When all DTX genes were considered in the aggregate, comparisons done between weevils feeding on different legume hosts had a significantly higher number of differentially expressed DTX genes. This also holds true when DTX genes are considered separately (Fig 1v). While the interaction of Host:Tissue was not significant for HD or IM genes, there was a significant interaction between tissue and host, and between tissue and functional gene group for DTX genes, suggesting that gene expression was influenced by the sampled tissue type for this class of genes (Table 1).

Heatmaps demonstrated that different host plant families required different degrees of species-specific attenuation for weevils, even if the hosts belong to the same group. For legume-legume comparisons, there was high expression intensity of DTX and IM genes but not of HD genes for adult tissues. For citrus-citrus comparisons, there were marked differences for expression intensity in HD and IM gene groups in both adult tissues, and in DTX genes only in immature tissues. For other-other (aster-aster) comparisons, there were strong expression intensity differences in IM and DTX genes in adult tissues (Fig 2i).

#### Switching host plants increases herbivore expression of host detection and detoxification genes in adults and pre-feeding immatures

The number of upregulated genes between weevils in the natal versus novel host plants was not significantly different for HD genes (Fig 1iv). When the number of upregulated DTX genes between the two conditions was analyzed as an aggregate, there were higher numbers of DTX genes upregulated in the switch condition. Additionally, some of the DTX genes showed significantly higher numbers of upregulated genes in the switch condition, such as *ABC transporters*, *cytochrome P450s*, and *glutathione S-transferases*. For IM genes, the number of upregulated genes was not significantly different between the two conditions (Figure 1iv). There was no identifiable interaction of sampled tissue type for any of the three target gene categories, as evidenced by non-significant Host:Tissue and Tissue:Functional Gene Group interactions (Table 1).

In the natal/novel heatmaps, red coloration indicates positive expression intensity for weevils in the switch condition, whereas blue coloration indicates positive expression intensity in the maintained condition. HD genes showed upregulation in the switched weevils (Fig 2i), suggesting that although the number of HD genes was not elevated in the switched weevils (Fig 1iv), the expression intensity of those DEGs was elevated. DTX gene expression intensities were also different, with all three tissue types registering higher median expression intensities in the switched condition. There were roughly equal levels of expression in IM genes for head and immature tissue between the switched vs. maintained weevils, indicating that similar numbers of different IM genes were expressed in both conditions at similar intensities. Abdominal tissue had a noticeably higher expression intensity than other tissue types in both the switch and maintain conditions (Fig 2i).

#### Common herbivore DEG patterns in the three targeted gene categories across different host plants and tissues

##### Expected and unexpected tissue-specific expression patterns in the three targeted gene categories

Expression of HD genes across weevil tissues presents a puzzling pattern. We expected expression to be more pronounced in adult head tissues, and that is true in *N. leucoloma* when feeding on legumes and when switching host plants (Fig 2i). However, we observed higher expression levels in abdomen tissues in conventionally grown citrus and when switching host plants. Interestingly, there was also strong expression of HD genes in immature tissues when parents fed on organically grown oranges.

Weevils feeding on different species of citrus host plants displayed higher expression levels of HD genes in all adult tissues relative to those feeding on different species of legume host plants (Fig 2ii). Interestingly, the number of DEGs was significantly larger in the legume to legume contrasts (Fig 1v).

As expected, expression of DTX genes was more prevalent in abdominal tissues when feeding on almost every host plant, including legumes, organically grown oranges and when switching host plants. However, we also saw strong expression of these genes in head tissues when feeding on non-legumes and on organically grown oranges. There was also strong expression of DTX genes in immature tissues when parents fed on legumes, organically grown oranges or on different species of conventionally grown citrus.

The expression of IM genes was expected to be equally prominent in both adult tissues (head and abdomen). That pattern is not seen in weevils feeding on legumes when contrasted with feeding on non-legumes, or on conventional citrus, where there is no measurable difference in expression in head tissues (Fig 2i). However, we saw generalized expression of IM genes in all adult tissues when feeding on conventional or organic oranges, on different species of citrus (Fig 2ii), or when switching to a novel host plant or maintaining the natal host plant (Fig.2i). We also observed widespread expression of IM genes to intermediate or high levels in immatures; this is true for almost all host plant contrasts (including within family contrasts) and experimental conditions.

##### Detoxification DEGs are more host-specific than host detection and immune defense DEGs

For host detection genes, the Legume vs. Other comparisons had the highest number of uniquely expressed genes, with 17 unique genes. Conventional vs. Organic and Switch vs. Maintain contained considerable overlaps with Legume vs. Other comparisons (31 and 37 genes, respectively) (Fig 3i). A set of 34 host detection genes were shared between all four comparison groups.

**Fig 3.**
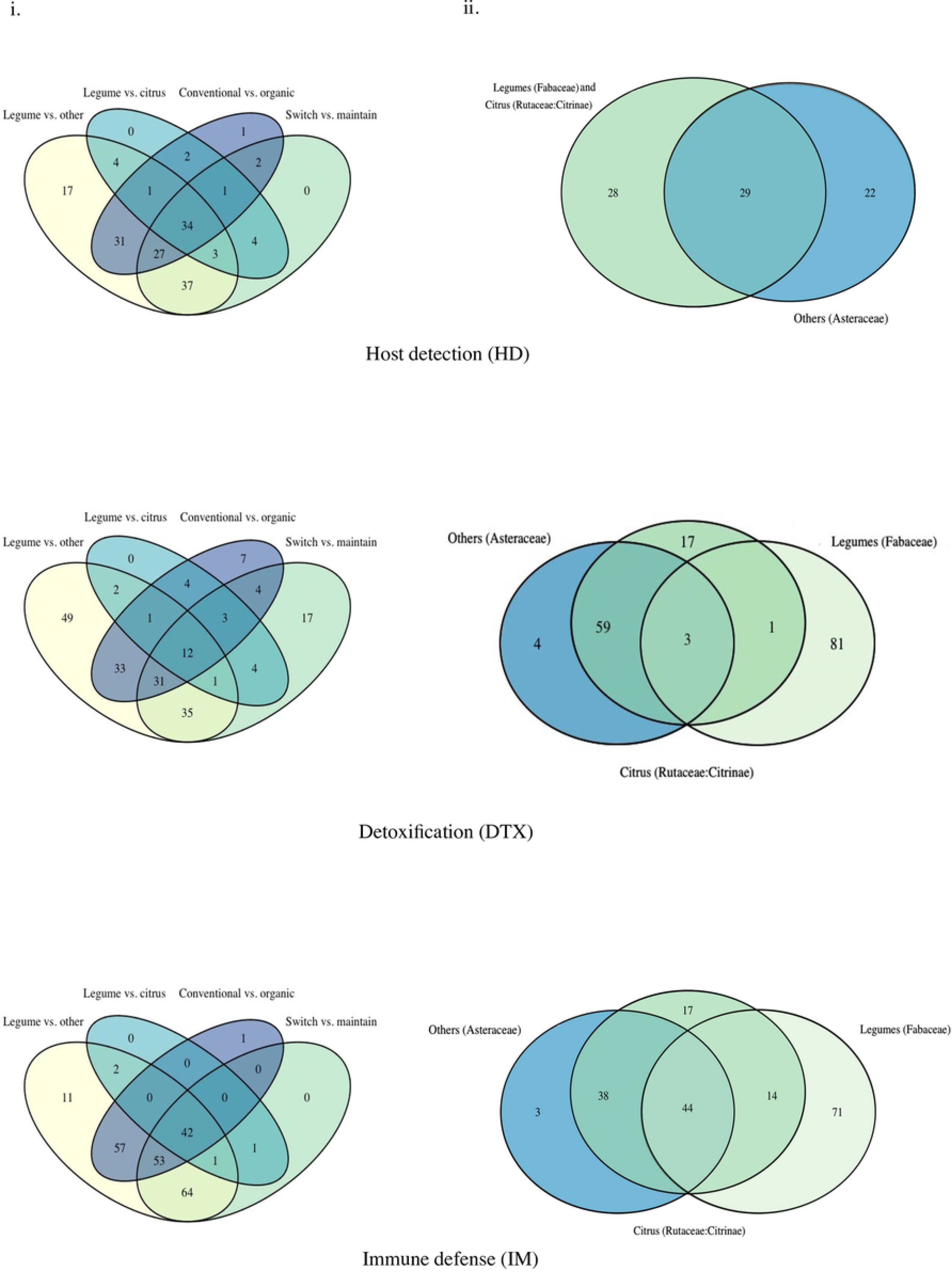
Number of unique and shared differentially expressed genes (DEGs) associated with host detection, host detoxification and immune defense between comparisons. Venn diagrams show overlaps or uniqueness in the identity of differentially expressed transcripts in either direction between comparisons. (i) four-way Venn diagrams including: Legume vs. Other, Legume vs. Citrus, Conventional vs Organic, and Switch vs. Maintain for host detection-related DEGs (HD), host detoxification-related DEGs (DTX) and immune defense-related DEGs (IM). (ii) three-way Venn diagrams including comparisons within the same host plant family: Citrus (Rutaceae: Citrinae); Legumes (Fabaceae); and Others (Asteraceae).

Venn diagrams for DEG identity in within host-family contrasts showed that there were no unique differentially expressed host detection genes for legume-legume (Fabaceae) nor citrus-citrus (Rutaceae) comparisons that were not shared between these two groups, which appears as a total overlap of 28 genes between the two families (Fig 3ii). There was an unexpectedly high number of host detection genes unique to the aster-aster host plant comparisons (Fig 3ii). Overall, there was a core set of 29 differentially expressed genes in common between all the included comparisons (Fig 3ii).

For detoxification genes, the Legume vs. Other comparisons had the highest number of uniquely expressed genes, with 49 unique genes, followed by Switch vs. Maintain comparisons, with 17 unique genes. Legume vs. Other and Switch vs. Maintain comparisons shared 35 detoxification genes. A set of 12 genes was shared between all four comparison groups (Fig 3i). As seen in other gene categories, the identities of many of the detoxification DEGs in weevils introduced to novel hosts overlap with those overexpressed in the legume contrasts.

In within-family comparisons, legume-legume contrasts had the most unique differentially expressed genes, with 81 genes falling into that category, followed by citrus-citrus comparisons with 17 (Fig 3ii). Interestingly, 59 genes were shared between other-other and citrus-citrus comparisons (Fig 3ii). There was a small core of detoxification genes shared by all comparisons (3 DEGs), contrasting with the high numbers of legume-specific DEGs (81 DEGs).

For immune defense genes, the Legume vs. Other comparisons again had the highest number of uniquely expressed genes, with 11 unique genes. There was a large number of shared genes between Legume vs. Other, Conventional vs. Organic and Switch vs. Maintain (57 and 64 genes, respectively). A core of 42 genes was shared between all four comparison groups (Fig 3i).

The comparisons of differentially expressed immune defense genes between weevils feeding on legumes had the most unique differentially expressed genes (77 DEGs), followed by citrus-citrus comparisons (17 DEGs); the Asteraceae (other) host plant comparisons had the fewest (3 DEGs) (Fig 3ii). There was also a core of 44 differentially expressed genes common to all comparisons (Fig 3ii).

### Exploration of transcriptome-wide expression patterns reveals common expression of GO terms between different host plants and treatments

To see if and how changes in global GO term enrichment were unique, or specific, to each comparison condition, Venn diagrams for GO identity uniqueness were constructed for both positive and negative enrichment directions. Positively enriched gene sets shared between comparisons of Legume vs. Other (enriched in legume-feeding weevils relative to weevils feeding on other hosts) and Legume vs. Citrus (enriched in legume-feeding weevils relative to citrus-feeding weevils) included GO terms for “ribosome”, “ribosomal construction”, “translation”, “chitin metabolic process”, and “chitin binding” (Fig 4i). Although one might expect that genes enriched in legumes would be entirely shared between these two comparison groups, because the Group 2 (other hosts vs. citrus hosts) is different, there were also different GO terms exclusive to the two comparison groups (8 and 7 exclusive GO terms for Legume vs. Other and Legume vs. Citrus, respectively) (Fig 4i). Interestingly, there was also an overlap of five enriched GO terms between the Legume vs. Citrus comparisons and the Switch vs. Maintain comparisons (enriched in weevils switched to feeding on a novel host relative to weevils maintained on their natal host). There were 11 GO terms unique to the Switch vs. Maintain comparison group, indicating enrichment exclusive to the switched weevils, which is a much larger set of unique GO terms compared to other contrasts (Fig 4i). Interestingly, all of the unique enriched GO terms in Switch vs. Maintain were found in adult tissues (Fig 2i).

**Fig 4:**
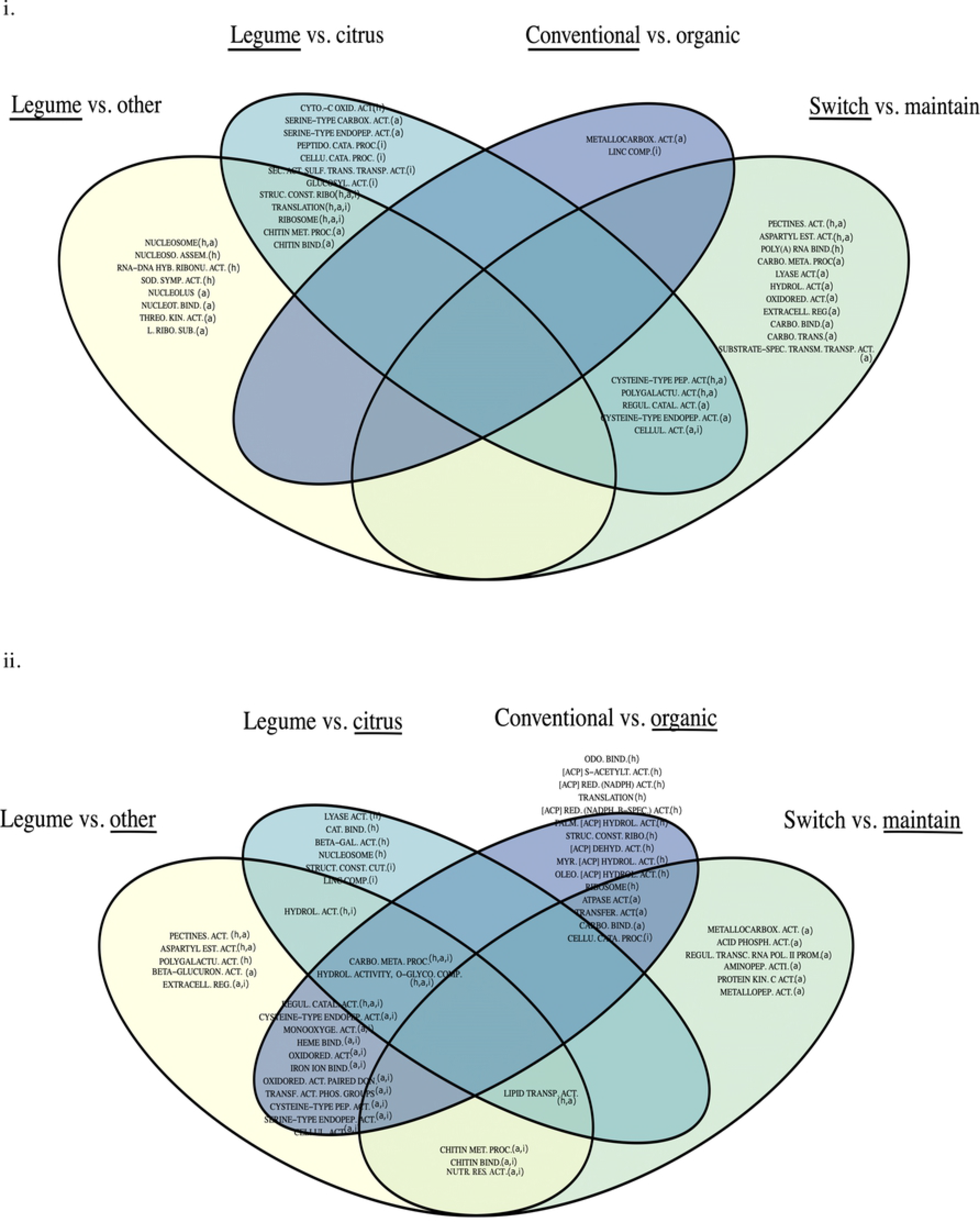
Exploration of generalized expression changes specific to particular host plants or experimental conditions. (i) Identity of upregulated Gene Ontology (GO) terms in G1; (ii) Identity of upregulated GO terms in G2, unique and/or shared between host plant contrasts and experimental conditions. Underline indicates the direction of the contrast, or which host plant group is overexpressing transcripts in that GO term. Parentheses after GO term descriptions contain the tissues where DE was found for each GO term (h: head, a: abdomen, and i: immature). Numerical identifiers for GO term abbreviations can be found in S3 Table.

Negatively enriched GO terms (indicating enrichment in Group 2, rather than Group 1, for a given comparison group) also contained overlapping enriched GO terms (Fig 4ii). Two GO terms, those for “carbohydrate metabolic process” and “hydrolase activity on O-glycosyl compounds”, were shared enriched terms between Legume vs. Other, Conventional vs. Organic, and Legume vs. Citrus (enriched in citrus-feeding weevils relative to legume-feeding weevils) comparisons (Fig 4ii). The more general GO term for “hydrolase activity” was an enriched shared term when either Citrus or Other were contrasted with Legumes (Legume vs. Citrus and Legume vs. Other)(Fig 4ii). Legume vs. Citrus comparisons contained a further 6 uniquely enriched GO terms (Fig 4ii). The Switch vs. Maintain contrast produced 7 uniquely enriched GO terms. Although it is difficult to identify why these terms are uniquely enriched in weevils maintained on their natal host, it is possible that transcripts that fall into these categories are instead underexpressed in weevils switching to a novel host plant.

Finally, Conventional vs. Organic comparisons contained the highest number of unique GO terms, with 15 terms enriched in weevils feeding on organically cultivated hosts. These included a highly interlinked set of acyl carrier proteins, which occurred in two different EnrichmentMaps with some modification (S2 Fig). The same network occurred again in immature tissue from citrus-citrus comparisons, as above but with the addition of two more GO terms (S2 Fig).

## Discussion

Several arthropod species show transcriptional plasticity in response to different host plant profiles (9, 46). In the same vein, this series of analyses sought to understand the processes of acclimation and adaptation to non-native host plants in two asexual *Naupactus* weevil species, as evidenced in their gene expression patterns.

### Taxing natal and novel host plants require highly specific transcriptional responses from herbivores

#### Legume and citrus host plants

Because legumes contain nitrogen-fixing rhizobia and generally have diverse repertoires of chemical defenses, there is a strong evolutionary pressure on legume-feeding herbivores to overcome these defenses in order to derive nitrogen for their own nutrition (25). This can result in a demonstrable preference for legume hosts (18), even though these legume species tend to require more energy-intensive herbivore responses to overcome the host’s defense response. Evidence of the extra cost imposed on legume-feeding weevils appears to be reflected in their gene expression profiles. The numbers of upregulated host detection genes, detoxification genes, and immune genes were significantly higher in legume-feeding weevils in both *N. cervinus* and *N. leucoloma* (Figs 1 and 2). This follows the prediction that both species invest more resources in detecting and dealing with secondary compounds of legumes, and that legumes elicit a larger immune response, possibly related to their associated rhizobia.

The identity of the overexpressed transcripts in legume-feeding weevils points to a legume-specific response. When examining both upregulated and downregulated host detection genes, detoxification genes, and immune genes, legume-feeding weevils had the highest number of unique differentially expressed transcripts (Fig 3i-ii), suggesting that the weevil’s transcriptional response pattern is highly specific to that host plant group. However, there were also strong overlaps in differentially expressed genes in comparisons between legume and other comparisons (Fig 3), suggesting that there are potentially shared mechanisms of responding to these particular host plants/growing conditions at the gene level. Previous work has hypothesized that host plant response specificity in herbivores may be exacerbated by the microbial communities specific to a host plant species, as ingesting microbes present on the leaf alters insect immunity (47). Adult *Naupactus* weevils feed on foliage and can encounter such leaf microbes.

#### Organically cultivated host plants

While the body of work contrasting transcriptional levels of defense compounds on conventional versus organic crops is not large, there is evidence that the production of some plant defensive compounds increases when plants are treated using organic rather than conventional approaches (48). Additionally, specific pathways related to RNA regulation and biotic stress have been found to be part of the variation in gene expression due to agricultural practices, with those pathways enhanced in organically fertilized or protected crops (49). In transcriptomes from weevils feeding on the same host species under different regimes of cultivation, there was a significantly higher quantity of upregulated host detection genes and specific categories of detoxification genes from weevils feeding on organically grown host plants (Fig 1b, Table 1). The expression intensity for differentially expressed immune genes was high across all three tissue types in both positive and negative directions (Fig 2). Taken as a whole, adult weevils feeding on organically raised hosts tend to elicit more upregulated genes in detoxification and host detection, with a slight trend in immune defense, supporting the hypothesis that organically cultivated host plants are associated with more differential gene regulation.

Even though organically raised host plants appear to challenge herbivores to a larger degree than their conventionally grown counterparts, the observed response in the three targeted gene categories is not unique to organically grown hosts. There was a notable overlap in the number of shared DEGs for host detection, detoxification and immune defense genes between Legume vs. Other and farming method comparisons (Fig 3). A greater degree of transcriptional plasticity and changes in genes associated with the metabolism of secondary compounds has been found as a response to exposure to stress in some aphids and other specialist insects (43, 50). The evolution of a conserved mechanism for both more toxic host plants and exposure to other forms of stress would be the least evolutionarily costly (51), and would be especially beneficial for this polyphagous species.

The pathway-level response to feeding on organically grown host plants included enriched GO terms in oxidation/reduction pathways, potentially linked to oxidative stress responses (S2 Fig). Transcripts involved in ribosome construction and translation are generally constitutively expressed, so it is interesting that our GO term analysis included these terms as significantly enriched. It may be the case that the enrichment of these terms points to an increase in translation of certain transcripts in response to xenobiotic compounds from resource-taxing host plants that require a change in weevil expression in basic metabolic pathways in order to clear these potentially life-threatening substances, as has been shown in *Helicoverpa armigera,* the polyphagous cotton bollworm (8).

Although the function of acyl carrier proteins in insect cells specifically is largely unknown (52), the uniquely citrus-specific enriched cluster of acyl carrier proteins found in organic-feeding weevils (Fig 4ii, S2iii Fig) is known to be linked to fatty acid biosynthesis and glycolytic pathways (53). This upregulation may indicate that the host plant defenses of organically treated oranges are more stressful for herbivores than those of conventionally treated oranges. Similar results have been proposed as a clear link between exposure to stress and increased transcriptional plasticity, including regulation of transcription and translation processes (43).

#### Short-term acclimation to a novel host plant

The important contribution of *cytochrome P450s* to the success of herbivore establishment on novel host plants has been previously documented in spider mites (28). In our experimental host plant switch, numbers of upregulated *ABC transporter, cytochrome P450,* and *glutathione S-transferase* genes were significantly higher in the switch condition (Fig 1b, Table 1).

A possible interpretation of the bidirectional nature of the expression of immune genes (Fig 2i) could be that the new host plant presents a new set of natural enemies, and as a herbivore feeds on a host where new natural enemies or parasites are present, immune genes associated with those pressures are regulated in one direction. Genes specific to the old host plant appear as regulated in the opposite direction, when in fact they may be simply maintained in weevils feeding on the old host relative to downregulation in weevils feeding on the new host. Support for the idea that herbivore detoxification and immune challenges are larger in newly colonized host plants is supported by the elevated herbivore diversity and load on native hosts relative to non-native hosts found in forty-seven different woody plant species (36).

While host detection and immune defense genes were entirely shared with other comparisons, a suite of 17 detoxification DEGs were uniquely specific to the Switch vs. Maintain contrasts (Fig 3). The length of host plant attenuation may explain the results described here; a host-plant specific set of detoxification genes may form the first line of short-term defense for a weevil introduced to a new host, as identified in spider mites challenged with new hosts of varying degrees of similarity in terms of secondary metabolites (29). On the other hand, long-term attenuation to a host plant may occur through host plant detection and immune pathways over longer timescales without exhibiting or requiring short-term specificity. It is possible that the investment needed to differentially regulate immune and host detection genes may come later as a long-term adjustment, whereas detoxification genes are differentially regulated early on to ensure survival on that new host. This is supported by other work indicating that a generalist’s short-term transcriptional response to a new host is detoxification-based, with the longer-term response including three times more differentially expressed genes across the genome (30).

Our results also present a set of 11 GO terms enriched exclusively in the switched weevils, providing a window into other pathways potentially involved in early acclimation to a new host plant. Some of these GO terms have been shown in other species to be highly variable and involved in stress responses to new environmental conditions (54). Other terms are implicated in the post-transcriptional regulation of mRNA maturation and export from the nucleus (55). This suggests that there are some upregulated GO terms related to responding to immediate environmental stress and the rapid adjustment of regulatory mechanisms that are enriched after a host plant transition. For parthenogenetic weevil species and other species with low genetic variation, an immediate response modulated by gene expression and epigenetic modification would be a useful way of acclimating quickly to new environmental conditions (56, 57). More generally, other arthropod studies that have examined new and old host plant adaptations in polyphagous insects have reported distinct transcriptional plasticity patterns during acclimation to such hosts (9, 29).

### Different modes of gene expression response: narrowly targeted vs. widespread

Even though our focus species have the potential to be polyphagous (58), individual weevil populations produce larvae that drop from the foliage to burrow into the soil to feed on the same host plant roots, which may result in the transgenerational extension of a specific host plant preference, regardless of polyphagous ability. Because of this dichotomy between potential and actual diet breadths, the expression of host-related genes in these weevils could take divergent modalities. Their patterns of gene expression may manifest as a widespread regulation of several common genes, as expected in a generalist species, or as a specific and targeted regulation of a few highly host-specific genes, as expected in a specialist species (8).

Citrus hosts appeared to elicit a narrow, targeted expression response of host detection genes in weevils feeding on different species of citrus hosts (Fig 2ii). One explanation for the targeted expression of host detection genes for citrus is the phylogenetic closeness of the citrus hosts examined here. Further research that corrects for this potentially confounding variable would be productive for more concretely identifying the source of this effect. However, this trend in highly specific, targeted expression for citrus hosts is replicated in other comparisons, and this pattern may be the result of acclimation to the unique chemical defenses of the host clade as well. Some research has found that the consequences of transfer to a new related host versus a new, distantly related host utilizes similar pathways (29). Following this idea of specialist, targeted expression, weevils feeding on a novel host plant increased expression intensity, but not number, of host detection and immune genes; such a targeted response was only observed in head and abdominal tissue, and may constitute the first signs of acclimation to the new host (Fig 2i).

It is important to note that the within-family comparisons involved two weevil species, with legume-feeding *N. leucoloma* compared against aster-feeding and citrus-feeding *N. cervinus*. Thus, for this particular set of comparisons, differences may be due to species biology rather than host plant attenuation. However, it is then interesting that host plant detection DEGs overlap entirely between two species feeding on legume and citrus host plants (Fig 3); either these species are very alike and the other results included in the host plant family analysis are credible, or the genes associated with host plant detection are highly conserved between species while the differentially expressed detoxification and immune defense genes have diverged. Host detection genes such as odorant-binding proteins are generally highly divergent between insect clades (Sun et al., 2018), suggesting that our results are probably due to genuine alterations in gene regulation patterns.

The contrasts between legume host plants show a pattern that appears to follow what would be expected for a generalist insect, with a high-intensity response involving large numbers of upregulated genes. This is particularly noticeable in detoxification genes, where the quantity of upregulated detoxification genes is significantly higher between weevils feeding on different legumes than in contrasts in the other two host plant groups (Fig 1b). Even though the modality of expression involving larger numbers of genes may appear like that of a generalist, the identity of the transcripts that are differentially expressed shows a large degree of specificity. Legume-feeding weevils had more total differentially expressed unique detoxification genes than either the citrus-feeding weevils or the aster-feeding (non-citrus, non-legume) weevils (Fig 3). This data supports the idea that observed differences in gene expression are highly dependent on the chemical characteristics of a specific host or plant family, or in this case, differences between members of the same host plant family. Legumes are not unique in eliciting specific defensive responses from herbivores; studies on Coleoptera and Lepidoptera feeding on Brassicaceae also respond specifically to the chemical defense profile of that host clade (60).

### Resource allocation and transgenerational gene expression effects

Transgenerational, host quality-dependent effects have been observed in insect herbivores before, as parental modulation of offspring phenotype can better adapt that progeny to different host plant qualities (61).

The intensities of gene expression for detoxification and immune defense genes were particularly interesting in comparing transcriptomes between immatures derived from legume-feeding versus citrus-feeding parents. In this case, the intensity of expression was strong in both upregulated and downregulated immune and detoxification genes (Fig 2i), suggesting that there are different sets of immune and detoxification genes that are differentially expressed between the offspring of legume-feeding weevils and citrus-feeding weevils. Overexpression of *cytochrome P450s* in larval stages has been reported in other citrus-feeding arthropods, such as the citrus red mite, but the role of this major detoxification enzyme has been linked to resistance to insecticides rather than to citrus-specific defenses (62). Legumes have the unique potential for rhizobia-mediated augmentation of host plant defenses (38, 63), and because of this, differences between immune gene regulation in legumes versus citrus were expected. However, this effect was only identifiable in immature tissue, which suggests that this pattern is potentially specific to this life stage.

From the GSEA results of weevils feeding on legumes versus non-legumes, it appears that GO terms associated with “ribosome assembly” and “nucleosomes” are enriched solely in adult tissues (Fig 4). The immature comparison yielded primarily downregulated gene sets, which may be the effect of resource allocation towards adult survival rather than host-priming of offspring (Fig 4). If the adult stage must dedicate its energy to surviving on a difficult host plant, previous research has suggested that this triggers the diversion of energetic resources away from reproduction and towards survival (64, 65), so that gene sets are less modulated in immatures from these adults relative to immatures from adults feeding on less well-defended host plants. In our experimental set-up, where immatures were processed before they were able to feed and therefore not yet exposed to the challenges presented by their host plants, the decreased parental investment in offspring priming would be more prominent. This would follow the above findings of generally higher numbers and combined expression indices of upregulated immune, detoxification, and host detection genes in adult weevils from both species that fed on legumes.

We see high expression intensity in immatures from parents feeding on organically raised host plants across all three gene groups, despite no significant difference in number of DEGs across these three categories, with the exception of HD genes (Fig 2). Because an organic host is not as difficult as a legume host, feeding on organic hosts allows for the parent to maintain any investment in reproduction and offspring host priming, rather than reallocating that energetic resource to immediate survival. The very low number of enriched gene sets in the immature comparison (S2iii Fig) from parents feeding on organically raised host plants could indicate that host plant cultivation may have less of an effect on regulation at the gene pathway level than host plant groups, as more enriched gene sets were observed in host plant group comparisons. Previous work has shown a highly specific gene response but common gene family response during a herbivore’s long-term acclimation to a particular host plant (29), and the low number of enriched pathways but significant difference of DEG number and expression intensity in this set of comparisons may support this finding.

A set of 11 enriched GO terms exclusive to those weevils that have fed on a new host plant are found only in adult tissues. It appears that transgenerational effects at the pathway level are not generalized, although at the gene level, a small effect was observed for detoxification and immune genes. This is surprising, because it would be reasonable to assume that any transgenerational transmission of the parent’s acclimated phenotype specific to a new host plant could help the offspring be better poised to face those same conditions. Transgenerational effects of environmental conditions have been recorded in asexual colembolan species and sexually reproducing grass moths (24, 66). However, it is also possible that a multi-pathway transgenerational enrichment may not be immediately needed, and that the more specific priming in the form of increased expression of specific detoxification and immune genes is enough of an advantage for offspring to survive.

### Closing Remarks

Our results have shown that the gene expression response of some *Naupactus* weevils can be specific to particular host plants, and that elements of that response can be transgenerational. Moreover, some host plant groups, such as legumes, appear to be more taxing to weevils as they elicit a complex gene expression response which is both strong in intensity and specific in identity. However, the weevil response to the secondary metabolites of taxing host plants shares many attributes (i.e., identity of upregulated transcript and enriched GO terms) with other stressful situations such as host plant cultivation conditions and/or a transition to a novel host, leading us to believe that there is an evolutionarily favorable core shared gene expression regime for responding to different types of stressful situations. Modulating gene expression in the absence of other avenues for phenotypic adaptation may be an important mechanism for successful host plant colonization for these introduced asexual insects.

## Experimental Procedures

### Weevil collection and rearing

Weevils were collected from Argentina in Buenos Aires and Entre Rios Provinces (7 localities) and within the United States in Georgia, Florida, Alabama (6 localities), and California (4 localities) (S1 Table). Adult weevils were maintained in temperature-controlled environmental rooms with 12:12 dark/light cycles at 24-28°C and 50% humidity (after Tarrant and McCoy, 1989) for a three-week acclimation period. Each set of weevils was fed their natal host plant. Weevil rearing boxes were checked daily for eggs, and juvenile specimens were separated, allowed to develop for 7-10 days, and frozen before active feeding on plant matter began. Adults were processed three weeks after the acclimation period. For a set of experimental host switch trials, individual adults were randomly assigned to continue consuming their natal host or to switch to a novel host plant after the three-week acclimation period and processed after an additional three weeks.

### Sample preparation, RNA extraction, and quality control

Adults and immature (larval) specimens were frozen and preserved in RNAlater (Invitrogen, Carlsbad, CA). RNA extraction was performed using the PureLink RNA Mini Kit (Ambion, Carlsbad, CA). To obtain enough RNA from adult tissues we extracted material from a pool of six weevils for the head and abdomen samples. To extract RNA from immature tissues, between 50 to 100 first instar larvae were pooled, also maintaining the parental host plant and locality of origin. RNA concentration and quality were assessed using a NanoDrop 2000 spectrophotometer (ThermoFisher Scientific, Carlsbad, CA) and a Qubit™ 4 fluorometer (Invitrogen, Carlsbad, CA). While a given tissue from a specimen pool representing a given locality was sequenced only once in this format, the differential expression analysis consists of comparisons of several of these pooled RNA samples.

RNA sequencing, transcriptome assembly, and initial GSEA results were completed by SeqMatic (Fremont, CA) from each of the 52 samples. The RNA-Seq libraries were compiled using the Illumina HiSeq 2500 platform (Illumina, San Diego, CA) and transcripts were assembled *de novo* using the R/Bioconductor (http://bioconductor.org) package `Trinity` (68) (data available in NCBI GEO, accession numbers….). Transcripts were mapped to identified/putative protein sequences in the UniProt database (http://uniprot.org), with the best hit used for transcript annotation and the assignment of gene ontology (GO) terms. Each sample transcriptome was aligned against an initial, arbitrary Trinity transcriptome assembly using the `bowtie` R package (69). The `RSEM` package (70) was used to calculate transcript and gene expression levels without the need for a reference genome.

Quality control measures were performed using FastQC (Illumina, San Diego, CA). Comparisons that included samples that were flagged during quality control analysis were not included (n=4), with the exception of two larval samples (Tul_onetwoC1I1 and Tul_threeC3I1), which were retained given that there were no replacements and the comparative paucity of immature samples. Only gene transcripts that had transcript counts of ≥10 in at least 1 sample were included for differential gene expression quantification. From 79,798 transcripts in *N. cervinus* samples, 54,366 genes were retained (68%); from 73,953 gene transcripts in *N. leucoloma* samples, 37,982 genes were retained (51%).

### Data processing and visualization

Gene expression levels were then assessed using FPKM and log_2_FC values. The fragments per kilobase of exon per million reads mapped (FPKM) is a normalized count value of the number of transcript fragments mapped onto a particular gene, corrected for the length of that gene and the sequencing depth. The log_2_-fold change in expression levels between the two groups of samples compared (log_2_FC) gives a relative measure of over- or under-expression for the sample groups being compared. For this analysis, a mapped gene was considered a differentially expressed gene (DEG) if the ΔFPKM was > 1 *and* the log_2_FC value was ≥ 1, indicating upregulation, or ≤ 1, indicating downregulation. All graphing was performed in RStudio v. 3.6.1 (see S4 Table) (71).

### Differential gene expression comparisons

Fifty-five individual samples were included in 48 pairwise comparisons (S2 Table). Although we did not sequence each sample multiple times, we obtained replicates by analyzing samples from similar tissues and host plants together, albeit from different localities. In particular, samples that fell into similar categories of tissue, host plant, plant family, plant farming method, or those maintained on the natal host plant or switched to a novel host plant were analyzed together in groups of varying sizes (1-7 samples per group), effectively acting as replicates. The expression levels between groups of samples were compared in a pairwise fashion. Both *N. cervinus* and *N. leucoloma* samples originating from native and introduced ranges were included when available but analyzed separately.

### Assessing upregulation in three targeted gene categories

To examine the role of IM, DTX, and HD gene regulation among host plant types and other conditions, composite violin/beeswarm DEG plots were constructed to visualize the number of differentially upregulated genes in categories of host detection genes (*odorant binding proteins, chemosensory proteins, gustatory proteins*), detoxification genes (*cytochrome P450s, glutathione S-transferases, glutathione peroxidases, ABC transporters, carboxylesterases, UDP-glycosyltransferases*) and immune defense genes (serine proteases/proteinases and serpins modulating the immune defense cascade, general immune response-related gene identities) in the pairwise comparisons used in each set. These were grouped by host plant, host plant family, plant farming method, or host switch condition (see Results), and broken into functional gene groups as defined above, as well as by tissue type. To analyze differential gene expression of a transcript with a certain functional annotation (i.e. odorant-binding protein, a host detection gene), the transcript must be present in both groups in a comparison. In cases where a transcript is annotated for a function of interest in one group (i.e. legume-feeding weevils in a Legume vs. Other comparison) but is not identified in the counterpart group (i.e. weevils feeding on other hosts in a Legume vs. Other comparison), the differential expression of that gene product is not able to be determined and is therefore excluded from the consequent violin plot, generating a different number of data points for each functional annotation within a plot. This does not exclude that comparison pair from being plotted for differential expression of other host detection genes (i.e. chemosensory proteins). The detoxification gene group was analyzed as both aggregate data, by summing the total number of upregulated genes in each condition, and separately by identity, by producing a violin plot that retained the gene’s functional identity information. To examine the potential interactions of sampled tissue type and functional gene group on the weevils’ expression response to different host plants, a rank-based nonparametric ANOVA was performed using the R package `Rfit` for each comparison group. If the interaction between a pair of variables was significant at □=0.05, the effect of the interaction was considered an influence on the distribution of the number of overexpressed genes in each comparison.

### Weighted expression heatmaps considering intensity of expression and number of differentially upregulated genes

Heatmaps were constructed to compare the weighted median intensity of expression (using log_2_FC values) in either direction for the three gene groups of interest. For each set of comparisons, DEGs falling into HD, DTX, or IM groups were separated, and log_2_FC ranges calculated separately for both positive and negative expression levels for each of those three gene groups. Comparisons that returned only one significantly upregulated or downregulated transcript in a gene group were excluded. These six expression range values (positive HD, negative HD, positive DTX, negative DTX, positive IM, and negative IM) were each split into five equal bins based on the range of expression values. This allowed the calculation of a median expression intensity, weighted by the number of genes in each bin, for both positive and negative expression in each of the three gene groups for each comparison. These weighted median expression intensities were then assembled into a heatmap and separated by tissue type for each comparison group.

Venn diagrams were employed to explore the number of shared or uniquely differentially expressed gene identities between comparisons. The same dataset used to build the plots for numbers of upregulated genes was used, separating by transcript identity between comparisons and retaining genes that are differentially expressed in either direction for HD, DTX, and IM genes. DEG specificity was visualized in a three-way or four-way Venn diagram, according to the comparison groups being tested.

### Exploration of global expression changes specific to particular host plants or experimental conditions

As *N. cervinus* and *N. leucoloma* are not model organisms, a preliminary investigation of global expression patterns associated with host plant use was also performed using Gene Set Enrichment Analysis (GSEA) (72). GSEA identifies functionally enriched pathways and/or families of genes for each comparison, producing a gene ontology (GO) term associated with each of these gene families/sets. Each of these sets are assigned an enrichment score, which indicates the degree to which the component genes of a gene set are overrepresented in that sample. This is normalized to ameliorate differences in gene set size, as some gene families are bigger or more researched than others, as well as differences in expression depth. Finally, a false discovery rate (FDR) is calculated to control for multiple testing and false positive errors.

### Network-based visualization of gene set enrichment patterns across all gene categories

To explore the relationships between these upregulated or downregulated enriched gene sets, a hierarchical clustering analysis of gene ontology terms was performed using the Cytoscape module EnrichmentMap (Cytoscape, v. 3.7.2) (73). Seventeen comparisons with the largest available sample sizes for each tissue class were selected to assemble EnrichmentMaps. Only gene sets with a false discovery rate (FDR) of < 0.05 and a log_2_FC > 1 were included to evaluate expression differences (74, 75). Cytoscape parameters were set so that *q* = 0.05, and the default connectivity level was employed. The gene set list files compiled from the initial transcriptome assembly for each species were used as references. This method of visualization allows for interpretation of overlaps between different GO terms/gene sets for a gene network-oriented analysis of regulation patterns at a global level, with stringent selective criteria. To examine differences and similarities in gene set enrichment between hosts and hypotheses, Venn diagrams were constructed, in this case separating GO terms by enrichment direction (positive or negative).

## Acknowledgements

We gratefully acknowledge the logistical support of Russ Mizzel from the NFREC-Quincy, FL, of Russell Ottens from the University of Georgia, and of Elizabeth Grafton-Cardwell and Joshua Reger from the Lindcove Research and Extension Center, University of California. We also greatly benefited from the guidance and field support of personnel at the USDA-ARS Appalachian Fruit Research Station, West Virginia, the USDA Southeastern Fruit and Tree Nut Research Laboratory in Georgia, the IFAS Extension, University of Florida in Homestead, Florida, the Florida Department of Agriculture and Consumer Services in Gainesville, Florida, and the Auburn University Gulf Coast and Chilton Research and Extension Centers in Alabama. Field assistance was provided by M. Koniger, J., M. and A. Rosado. J. Gums and C. Hou provided laboratory assistance.

## Supporting Information

**S1 Fig. Individual heat maps displaying expression intensity for significantly up- and downregulated host detection (HD), host detoxification (DTX) and immune defense (IM) genes, including all tissue types for *N. cervinus* weevils feeding on different host plants or in different experimental conditions.** (i) host-specific expression between weevils feeding on a) Legumes vs. Other (for *N. cervinus* and *N. leucoloma*), b) Legumes vs. Citrus, c) Conventional vs Organic orange hosts and d) Switch vs Maintain. (ii) contrasts between expression levels while feeding on host plants from the same family Citrus vs. Citrus (Rutaceae:Citrinae), Legume vs Legume (Fabaceae) and Other vs Other (Asteraceae). Shades of red indicate upregulation in Group 1 while shades of blue indicate upregulation in Group 2. Gray indicates that a median differential expression value was not calculated due to a low DEG count.

**S2 Fig. EnrichmentMaps displaying differentially coexpressed gene sets, as determined by GSEA products**. Gene Ontology (GO) term coloration in red indicates upregulation in G1, whereas blue coloration indicates upregulation in G2; the host plant listed first always corresponds to G1. (i) Weevil transcriptome comparisons while feeding on Legume vs. Other host plants: a) *N. cervinus* head tissue comparison (Comparison C25i/n). b) *N. cervinus* abdomen tissue comparison (Comparison C24i/n). c) *N. cervinus* immature tissue comparison (Comparison C56i). (ii) Weevil transcriptome comparisons while feeding on Legume vs. Citrus host plants: a) *N. cervinus* abdomen tissue comparison (Comparison C67i). b) *N. cervinus* head tissue comparison (Comparison C66i). (iii) Weevil transcriptome comparisons while feeding on Conventional vs. Organic host plants: a) *N. cervinus* abdomen tissue comparison (Comparison C39i). b) *N. cervinus* head tissue comparison (Comparison C38i). c) *N. cervinus* citrus-citrus immature tissue comparison (Comparisons C52i and C46i). (iv) Weevil transcriptome comparisons while feeding on host plants within the same host-plant family: a) *N. cervinus* citrus-citrus abdomen tissue comparison, (Comparisons C51i and C45i). b) *N. cervinus* citrus-citrus head tissue comparison (Comparisons C50i and C44i). c) *N. cervinus* citrus-citrus immature tissue comparison (Comparisons C52i and C46i). Left hemisphere represents an organic orange vs. rough lemon comparison, whereas the right hemisphere represents a conventional orange vs. rough lemon comparison. d) *N. cervinus* aster-aster abdomen tissue comparison (Comparison C26i/n). e) *N. cervinus* head tissue comparison (Comparison C27i/n). (v) Switched vs. Maintained host plant weevil transcriptome comparisons: a) *N. cervinus* abdomen tissue comparison (Comparison C70i). b) *N. cervinus* head tissue comparison (Comparison C69i).

**S1 Table. List of collection records and samples organized by area.** General area indicates if the weevils were gathered from the introduced (INT) or native (NAT) range. Locality name and Coordinates provide locality details with location and state or provice codes or names. Host plant indicates the plants where weevils were collected from and were maintained in those hosts while in the lab. When localities had multipe hosts, those are numbered and included in the lab sample code. Lab sample codes include locality code with species designation (C and L), host number in that locality (some localities yielded samples from multiple host plants), tissue (A: head, B: abdomen, I: immature) and preparation number. For samples involved in the switch experiment, numbers in parentheses after the host label indicate switched to a new host plant (1) or maintained in the natal host plant (2) (for example: “Quin71C1(2)A1” denotes the first RNA preparation of head tissue from *N. cervinus* collected in FL on the one host present in that locality and maintained in that natal host). Comparisons indicates in which comparison groups those samples were included. Details of the comparisons are provided in Supplementary Table 2.

**S2 Table. Details of each group of contrasts used for differential expression analysis organized by prediction.** Comparison names include species label: C = *N. cervinus*; L = *N. leucoloma;* number within that species and range: i = both sample groups originated from the introduced range; i/n = sample groups are from different ranges, one from introduced range and one from the native range; n = both sample groups originated from the native range. Comparison details include Species name, host plant, host plant groups, or experimental condition for each sample group. Comparison groups display lab sample codes as detailed in S1 Table, which also includes the geographic origin of each sample.

**S3 Table. Summary of significantly enriched GO terms derived from Enrichment Maps for each comparison and tissue.** Significantly enriched GO terms are displayed by hypothesis, species and tissue. Within each contrast, the numbers of significantly enriched GO terms and the direction of enrichment are indicated together with the number of connections between GO terms in each direction produced by Cytoscape.

**S4 Table. Summary of R packages used in this study.** List of R software and packages including the sources for each module.

